# Subcellular localization of the J-protein Sis1 regulates the heat shock response

**DOI:** 10.1101/2020.04.02.022491

**Authors:** Zoe A. Feder, Asif Ali, Abhyudai Singh, Joanna Krakowiak, Xu Zheng, Vytas P. Bindokas, Donald Wolfgeher, Stephen J. Kron, David Pincus

## Abstract

Cells exposed to heat shock induce a conserved gene expression program – the heat shock response (HSR) – encoding chaperones like Hsp70 and other protein homeostasis (proteostasis) factors. Heat shock also triggers proteostasis factors to form subcellular quality control bodies, but the relationship between these spatial structures and the HSR is unclear. Here we show that localization of the J-protein Sis1 – a co-chaperone for Hsp70 – controls HSR activation in yeast. Under nonstress conditions, Sis1 is concentrated in the nucleoplasm where it promotes Hsp70 binding to the transcription factor Hsf1, repressing the HSR. Upon heat shock, Sis1 forms an interconnected network with other proteostasis factors that spans the nucleolus and the surface of the cortical ER. We propose that localization of Sis1 to this network directs Hsp70 activity away from Hsf1 in the nucleoplasm, leaving Hsf1 free to induce the HSR. In this manner, Sis1 couples HSR activation to the spatial organization of the proteostasis network.

**One sentence summary:** Localization of the J-protein Sis1 to a subcellular network of proteostasis factors activates the heat shock response.

## INTRODUCTION

Protein homeostasis (proteostasis) describes a cellular state in which protein synthesis, folding and degradation are balanced (*1*). When a cell has achieved proteostasis, molecular chaperones, sequestrases and degradation factors – collectively referred to as the proteostasis network (PN) – are expressed at sufficient levels such that nascent, misfolded and aberrant proteins are efficiently folded, triaged and degraded and the integrity of the proteome is maintained (*2-4*). Environmental fluctuations such as changes in temperature, nutrient availability and signals from other cells can increase the burden on the PN and overwhelm its capacity. In addition to environmental sources, neurodegenerative disorders such as Parkinson’s disease and amyotrophic lateral sclerosis (ALS) are characterized by protein aggregates and linked to deficits in PN components (*5-7*). By contrast, aggressive human cancers have been shown to usurp the PN to support malignant growth in the presence of high mutational loads (*8-10*). Thus, modulation of PN expression has been proposed as a therapeutic avenue to treat both cancer and neurodegenerative diseases (*11, 12*).

In eukaryotes, heat shock factor 1 (Hsf1) regulates a transcriptional program known as the heat shock response (HSR) that activates expression of chaperones like Hsp70 and Hsp90 along with a suite of other of PN genes (*13, 14*). The prevailing model for Hsf1 activation is a “chaperone titration model” in which the accumulation of chaperone clients that result from an overtaxed PN outcompete Hsf1 for access to chaperones, leaving Hsf1 free to induce the HSR (*15, 16*). Several different chaperones have been implicated in Hsf1 repression including Hsp90, Hsp70, J-proteins and TRiC/CCT (*14, 17-21*). Recently, multiple studies have pointed to Hsp70 as the primary and direct negative regulator of Hsf1 activity (*15, 16, 22*). Since Hsp70 is a major transcriptional target of Hsf1, these two proteins form a negative feedback loop that controls the dynamics of HSR induction (*23*). Although Hsp70 has emerged as a key repressor of Hsf1, roles for the other chaperones have not been ruled out.

While broadly consistent with existing data, the Hsp70 titration model does not account for the spatial organization of the cell in general or the protein quality control machinery in particular. First, the clients that titrate Hsp70 away upon heat shock are thought to be primarily nascent proteins emerging from ribosomes in the in the cytosol (*15*). Yet, Hsf1 activates target gene transcription in the nucleus in less than a minute following heat shock (*24*). How do cytosolic clients titrate away nuclear Hsp70 so quickly, especially given that the molecular ratio of Hsp70:Hsf1 in yeast cells exceeds 1000:1 (*25*)? Second, misfolded reporter proteins and components of the PN including sequestrases and disaggregases localize to specific subcellular sites in the cytosol (*26-28*). How does the stress-dependent spatial reorganization of the PN connect to HSR activation? I.e., how does the cell biological response coordinate with Hsf1 to modulate the transcriptional response?

In this study, we combine chemical genetics, single cell reporters, transcriptomics, proteomics, mathematical modeling and 3D live cell imaging to investigate the connection between the PN and the HSR in budding yeast. We identify the conserved J-protein Sis1 as a key factor required for Hsp70-mediated repression of Hsf1. J-proteins deliver clients to Hsp70 and activate Hsp70 to bind the clients with high affinity (*29*). Under nonstress conditions, Sis1 localizes to the nucleoplasm where it targets Hsp70 to bind and repress Hsf1. Upon heat shock, Sis1 re-localizes to the nucleolar periphery and the cytosolic face of the endoplasmic reticulum (ER) where it forms a semi-contiguous meshwork with other PN factors. Our data support a model in which, during heat shock, Sis1 targets Hsp70 activity to: 1) unincorporated ribosomal proteins condensed on the surface of the nucleolus and 2) ribosome-nascent chain complexes on the ER. This depletes Hsp70 activity from the nucleoplasm, leaving Hsf1 free to activate the HSR. The rapid reorganization of Sis1 into a two-dimensional network on the nucleolus and ER thus provides a mechanism for near-instantaneous activation of Hsf1 upon heat shock. In this manner, Sis1 localization dynamics relay the state of the proteostasis network to Hsf1 to modulate the HSR accordingly.

## RESULTS

### Chaperone anchor-away reveals that nuclear Sis1 represses Hsf1 activity

Multiple chaperones have been implicated in regulating Hsf1 to repress the heat shock response including Hsp90, Hsp70 and J-proteins (*17, 18, 20*). To determine which of these function as genetic repressors of Hsf1 in our yeast background, we deleted the genes encoding these chaperones in a strain bearing a fluorescent reporter of Hsf1 transcriptional activity and measured the reporter levels by flow cytometry under nonstress conditions (Table S1). The reporter consists of four repeats of the Hsf1 DNA binding site known as the heat shock element (HSE) driving yellow fluorescent protein (YFP) (Figure 1A, inset) (*16, 23, 30*). Consistent with previous high throughput experiments (*20*), HSE-YFP levels were significantly increased in cells lacking the highly expressed Hsp70 paralogs (*ssa1*Δ and *ssa2*Δ) but not in cells lacking the stress-inducible paralogs (*ssa3*Δ and *ssa4*Δ). Likewise, cells lacking the highly expressed Hsp90 paralog (*hsc82*Δ) showed increased HSE-YFP levels, but cells lacking the stress inducible paralog (*hsp82*Δ) had a normal response. We also observed increased reporter levels in cells lacking a J-protein (*ydj1*Δ) (Figure 1A). All mutants with elevated reporter expression retained the ability to further induce the HSE-YFP reporter upon heat shock (Figure S1A). The J-protein Sis1 is required for viability (*31*), so we could not evaluate its role via knockout. Double knockout of Hsp70 paralogs (*ssa1*Δ *ssa2*Δ) resulted in synergistic activation of the HSE-YFP reporter to over 20-fold above wild type levels. Triple mutants *ssa1*Δ *ssa2*Δ *ssa3*Δ and *ssa1*Δ *ssa2*Δ *ssa4*Δ were inviable (Figure S1A). Likewise, a double knockout of the Hsp90 paralogs (*hsc82*Δ *hsp82*Δ) was inviable (*32*). These genetic results suggested that Ssa1/2, Hsc82 and Ydj1 could all be Hsf1 repressors.

**Figure 1.**
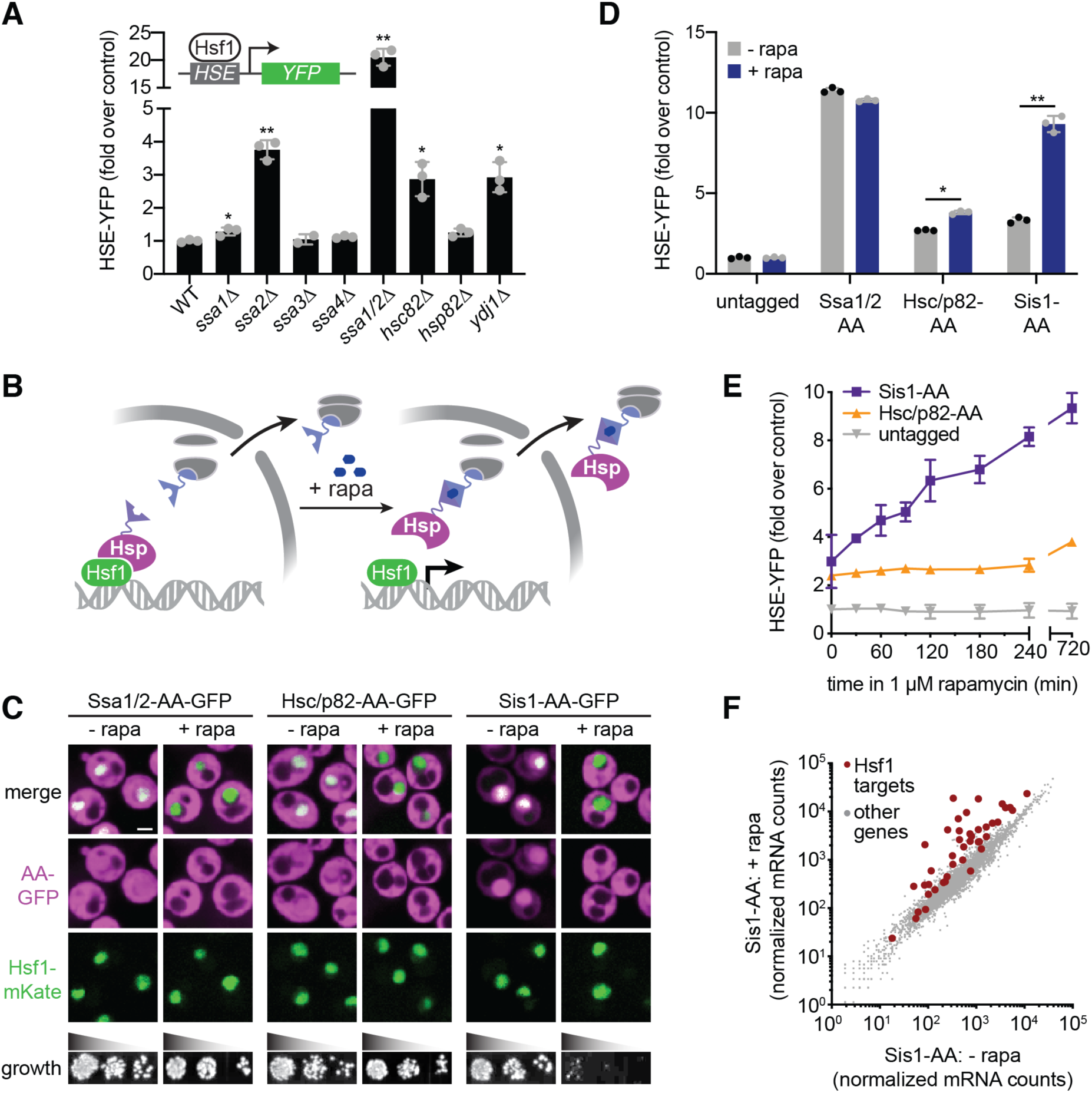
Nuclear depletion of Sis1 rapidly and specifically activates Hsf1. **A)** Heat shock element (HSE)-YFP reporter assay for Hsf1 activity. Cells were grown the absence of stress and YFP levels were measured by flow cytometry and normalized to wild type. Three biological replicates are shown, along with the mean and standard deviation. Inset: Cartoon of HSE-YFP reporter construct. **B)** Cartoon of the chaperone anchor away approach. Upon addition of rapamycin, putative chaperones are tethered to a ribosomal protein and depleted from the nucleus. **C)** Spinning disc confocal images of cells expressing Hsf1-mKate and GFP-tagged versions of Hsp70 (Ssa1/2), Hsp90 (Hsc/p82) and Sis1. Cells were treated with 1 µM rapamycin for 30 minutes before imaging. Scale bar is 2 µm. Lower panel: Indicated strains were serially diluted, spotted and grown at 30°C for 36 hours on YPD -/+ 1 µM rapamycin. **D)** HSE-YFP reporter assay of anchor away strains in the presence and absence of rapamycin normalized to the untagged anchor away parent strain. Cells were treated with 1 µM rapamycin for 8 hours before measuring the reporter. **E)** Time course of HSE-YFP levels in the Sis1 and Hsp90 anchor away strains compared to the untagged parent following addition of 1 µM rapamycin. **F)** RNA-seq analysis of the Sis1 anchor away strain in the absence and presence of 1 µM rapamycin for 30 minutes. The 42 known Hsf1 target genes are shown in red.

The deletion strains neither allowed us to assess the acute effects of chaperone depletion nor to determine the contribution of essential genes. To conditionally re-localize chaperones away from Hsf1, we used the anchor away (AA) approach to enable rapamycin-inducible depletion from the nucleus (Figure 1B) (*33*). In rapamycin-resistant strains, we AA-tagged Ssa1, Ssa2, Hsc82, Hsp82, Ydj1 and Sis1 (Figure S1B). We also doubly tagged Ssa1 and Ssa2 in one strain (Ssa1/2-AA) and Hsc82 and Hsp82 (Hsc/p82-AA) in another strain. GFP-tagged versions of Ssa1/2-AA and Hsc/p82-AA all localized equally throughout the cytosol and nucleus, while Sis1-AA was concentrated in the nucleus (Figure 1C). We found that rapamycin selectively depleted Ssa1/2-AA, Hsc/p82-AA and Sis1-AA from the nucleus while leaving Hsf1-mKate in the nucleus (Figure 1C). Rapamycin had no discernable effect on proliferation in Ssa1/2-AA and Hsc/p82-AA, but it inhibited cell growth in Sis1-AA cells (Figure 1C).

To assay the effects of anchoring away the chaperones on Hsf1 activity, we measured the HSE-YFP reporter. In the Ssa1/2-AA strain, we observed more than a 10-fold increase in HSE-YFP levels relative to the untagged parent strain in the absence of rapamycin, but we found no further increase upon rapamycin addition (Figure 1D). Indeed, we determined that tagging Ssa1/2 at either terminus impaired function, rendering anchor away of Hsp70 unmeaningful (Figure S1B). Hsc/p82-AA cells remained viable, indicating that the tagged versions retained essential Hsp90 functionality. The HSE-YFP reporter was modestly increased Hsc/p82-AA cells relative to an untagged strain in the absence of rapamycin. Addition of rapamycin resulted in a small but significant further increase in HSE-YFP levels (Figure 1D). However, it took multiple hours for nuclear depletion of Hsc/p82-AA to increase HSE-YFP levels (Figure 1E). Sis1-AA also displayed increased basal HSE-YFP signal. Yet, addition of rapamycin strongly induced the HSE-YFP reporter in the Sis1-AA strain (Figure 1D). Anchor away of Sis1-AA led to an immediate and sustained increase in HSE-YFP signal (Figure 1E). The magnitude and immediacy of HSE-YFP induction following Sis1 anchor away suggested that Sis1 plays a major role in Hsf1 repression under nonstress conditions.

If Sis1 is required for Hsf1 inactivation, then its nuclear depletion should activate Hsf1 without causing protein homeostasis (proteostasis) collapse, and it should induce the full Hsf1 target gene regulon without activating other stress pathways. As a proxy for global proteostasis, we monitored the localization of Hsp104, a disaggregase known to form foci thought to mark protein aggregates (*34*). Heat shock resulted in significant accumulation of Hsp104-mKate foci. By contrast, anchoring away Sis1 did not alter Hsp104-mKate localization, suggesting that proteostasis remained largely intact (Figure S2). To measure the transcriptional changes triggered by anchor away of Sis1, we analyzed global mRNA expression by mRNA deep sequencing (RNA-seq) following either rapamycin addition or heat shock in the Sis1-AA strain. Heat shock resulted in induction of the full Hsf1 regulon (*35*), but also induced and repressed hundreds of other genes (Figure S3A) (*36*). By contrast, Sis1 anchor away specifically induced the Hsf1 regulon without other changes to the transcriptome (Figure 1F, S3B). Moreover, while Hsf1 target mRNA expression peaked after 15 minutes of heat shock and subsequently declined, Sis1-AA resulted in sustained expression of the full Hsf1 regulon, albeit to a lower magnitude than heat shock (Figure S3C-F).

### Nuclear Sis1 maintains the interaction between Hsp70 and Hsf1

We next tested if Hsf1 forms a protein complex with Sis1. We performed anti-FLAG immuno-precipitation (IP) of Hsf1-3xFLAG from unstressed cells and from cells that had been heat shocked for 60 minutes to induce high-level expression of Sis1. While we efficiently pulled down Hsf1, mass spectrometry (MS) analysis failed to identify Sis1 as an interactor in either condition (Table S2). Only Ssa1/2/3/4 (Hsp70 paralogs) associated specifically with Hsf1, consistent with our previous results (Figure 2A) (*16*).

**Figure 2.**
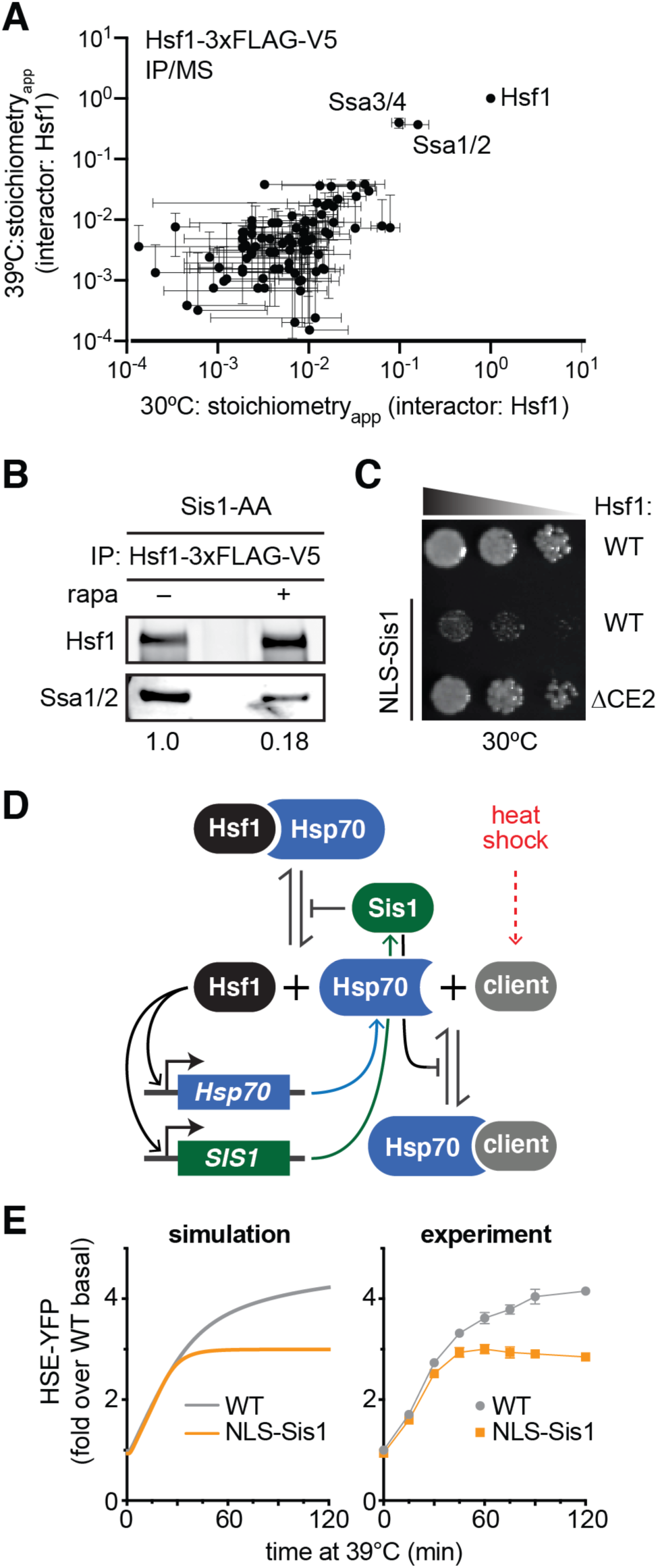
Sis1 promotes interaction between Hsf1 and Hsp70. **A)** Anti-FLAG Immunopreciptation (IP) of Hsf1-3xFLAG-V5 followed by mass spectrometry at 30°C and at 39°C for 60 minutes. IPs were performed in biological triplicate. Levels of interacting proteins were plotted relative to the amount of Hsf1 measured in each replicate to generate an “apparent stoichiometry” value. Mean and standard deviation of the replicates are shown. **B)** Anti-FLAG/V5 serial IP of Hsf1-3xFLAG-V5 followed by anti-FLAG and anti-Ssa1/2 westerns. Cells were treated in the absence and presence of 60 minutes of 1 µM rapamycin treatment to anchor away Sis1. Values are ratios of Ssa1/2:Hsf1 in each lane (average of three replicates). **C)** Dilution series spot assay. Ectopic expression of NLS-Sis1 impairs cells growth; deletion of the CE2 region of Hsf1 rescues growth in NLS-Sis1 cells. Cells were grown for 36 hours on YPD + 20 nM estradiol to induce NLS-Sis1. **D)** Schematic of the mathematical model of the Hsf1 regulation by the Sis1 and Hsp70. During heat shock, Hsp70 clients accumulate and compete for Sis1 and Hsp70 and release Hsf1. Active Hsf1 then induces expression of Sis1 and Hsp70. **E)** Simulations of the mathematical model and corresponding experiments of heat shock time courses of HSE-YFP reporter strains with and without ectopic expression of NLS-Sis1. NLS-Sis1 was induced with 20 nM estradiol for 1 hour prior to the time course.

As a J-protein, Sis1 is thought to deliver clients to Hsp70 and activate high affinity Hsp70 binding (*29*). Since we found no evidence that Sis1 forms a stable complex with Hsf1, we hypothesized that Sis1 promotes the interaction between Hsf1 and Hsp70 but does not remain part of the mature complex. To test this, we performed IPs of Hsf1-3xFLAG and blotted for Hsp70 (Ssa1/2) in a Sis1-AA strain in the presence and absence of rapamycin. Anchoring away Sis1 resulted in greater than a 5-fold decrease in the amount of Ssa1/2 that co-precipitated with Hsf1-3xFLAG (Figure 2B). Thus, in the absence of nuclear Sis1, the interaction between Hsp70 and Hsf1 is reduced.

As an additional piece of evidence that Sis1 mediates its repressive effect on Hsf1 by promoting Hsp70 binding, we identified a genetic interaction relating Sis1 to Hsf1 and Hsp70. We found that ectopic expression of Sis1 fused to a nuclear localization signal (NLS-Sis1) impairs growth in wild type cells under nonstress conditions. However, this growth phenotype is suppressed in cells expressing Hsf1ΔCE2 as the only copy of Hsf1. Hsf1ΔCE2 lacks a binding site for Hsp70 that represses Hsf1 activity (Figure 2C) (*22, 23*). Thus, relieving Hsf1 repression by Hsp70 is sufficient to rescue growth in cells with too much Sis1 in the nucleus. This result implies that overexpression of Sis1 in the nucleus leads to hyper-repression of Hsf1 by Hsp70, to the point that Hsf1 ceases to perform its essential basal transcriptional function. Without the CE2 binding site for Hsp70, NLS-Sis1 cannot completely inactivate Hsf1. Together, these biochemical and genetic data support a role for Sis1 in repressing Hsf1 via Hsp70.

### Mathematical modeling of the Sis1-Hsp70-Hsf1 regulatory circuit

To formalize the role of Sis1 in the Hsf1 regulatory circuit, we generated a mathematical model of the heat shock response. Like our previous versions (*16, 23*), the model is based on a negative feedback loop in which Hsf1 activates Hsp70 expression, and Hsp70 represses Hsf1 activity. Heat shock generates chaperone clients that accumulate and titrate away Hsp70, releasing active Hsf1 (Figure 2D). In this version of the model, Sis1 acts to increase Hsp70 binding affinity to both Hsf1 and other clients, and Sis1 expression is also driven by free Hsf1 (Figure 2D). The output of the model is expression of HSE-YFP, enabling comparison to the cellular response (see methods).

We first confirmed that the model was able to recapitulate the dynamics of the heat shock response by comparing a simulation of the HSE-YFP reporter over a heat shock time course in wild type cells to experimental data (Figure 2E). Next, we tested if the model was capable of capturing the effect of overexpression of Sis1. The model only simulates the cell nucleus, so overexpression of Sis1 in the model is equivalent to an increase in nuclear Sis1 in cells. The model predicted that increased expression of Sis1 in the nucleus should reduce the maximum HSE-YFP output over a heat shock time course and attenuate the response faster than WT (Figure 2E). To test this prediction experimentally, we generated a strain with the HSE-YFP reporter in which we could induce ectopic expression of Sis1 with an appended nuclear localization signal (NLS-Sis1) (*37*). We performed a heat shock time course from 25°C to 39°C for two hours in cells with and without induction of ectopic NLS-Sis1. In agreement with the model, cells expressing NLS-Sis1 failed to reach the maximal HSE-YFP level achieved by WT and deactivated more quickly (Figure 2E). These results demonstrate that a model based on the Sis1-Hsp70-Hsf1 regulatory axis is consistent with experimental data and such a model can quantitatively recapitulate the effects of perturbations to Sis1.

### Sis1 interacts with chaperones, proteasomes, ribosomes and RQC during heat shock

Our chemical genetic experiments and mathematical modeling suggest that Sis1 is a key regulator of the HSR, but provide little insight into the physiological role of Sis1 during heat shock. To identify endogenous proteins that interact with Sis1, we performed IP-MS of Sis1-3xFLAG following a 15-minute heat shock. We identified 192 proteins with >99% confidence in three biological replicates (Table S3), and Sis1 was both the most abundant and most significantly enriched protein in the set. In Figure 3A, we plot the significance with which each of these proteins was enriched over an untagged control as a function of the relative abundance of each protein with respect to Sis1 (“stoichiometry_app_”). Gene ontology analysis of these proteins revealed functional enrichment for ribosomal proteins and biogenesis factors, stress granule components, proteasome subunits, chaperones, glycolytic enzymes and the Paf transcription elongation complex (Figure 3B). In addition to these categories, we also observed interactions with nucleolar factors, members of the ribosome-associated quality control complex (RQC) and the ER structural protein Rtn1 (Figure 3A, C). The heat shock-dependent Sis1 interaction network thus appears to link major cellular chaperone systems – Hsp70, Hsp90 and TRiC/CCT – with other major quality control systems including the RQC, stress granules, and the proteasome. The mix of nucleolar, cytosolic and ER factors among the Sis1 interactors suggests that Sis1 localizes to a complex subcellular network during heat shock.

**Figure 3.**
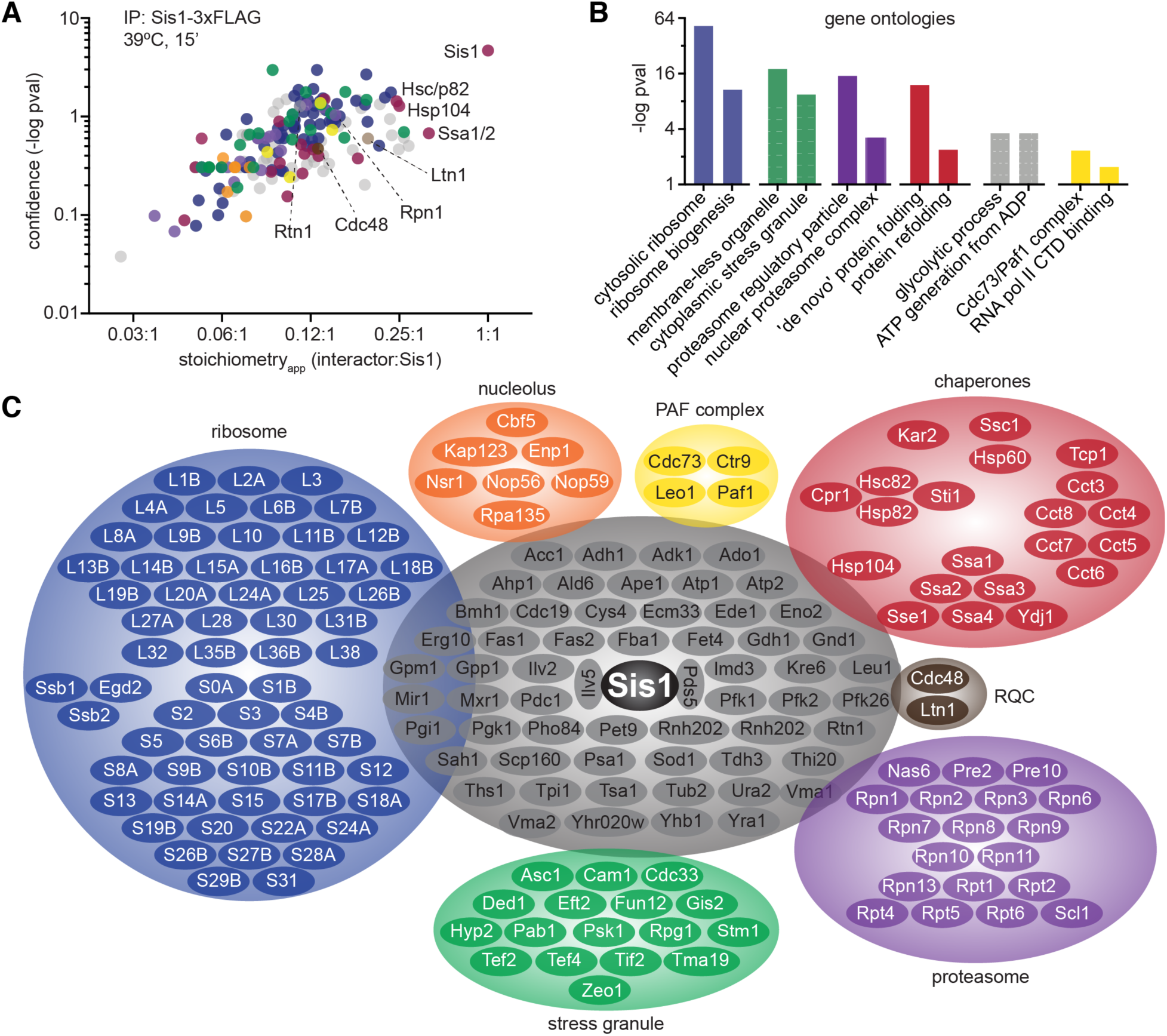
Sis1 localizes to a peri-nucleolar ring and cytosolic foci during heat shock. **A)** Live cell heat shock time course of cells expressing endogenously-tagged Sis1-YFP and Hsp104-mKate. Cells were imaged by spinning disc confocal microscopy. Scale bar is 2 µm. **B)** Cells expressing endogenously tagged Sis1-YFP and the nucleolar marker Cfi1-mKate imaged under nonstress and heat shock conditions. Scale bar is 2 µm. **C)** Scans of fluorescence signal for Sis1-YFP (black) and Cfi1-mKate (purple) along the dashed lines in the merged images in (B). **D)** Cells expressing endogenously-tagged Sis1-YFP and Hsp104-mKate following 10 minutes of heat shock after either no pretreatment or pretreatment with cycloheximide (CHX) to arrest protein synthesis. Scale bar is 2 µm. **E)** Quantification of the fraction of cells with a Sis1 nuclear ring (black) and Hsp104 foci (purple) after either no pretreatment or pretreatment with cycloheximide (CHX). Experiments performed in triplicate with >50 cells in each replicate. **F)** Live cells expressing endogenously tagged Sis1-mScarlet, Hsf1-YFP and Hsp104-BFP were imaged in a lattice light sheet microscrope under nonstress conditions and following heat shock at 39°C for 15 minutes. Scale bar is 2 µm. **G)** Scans of fluorescence signal for Sis1-mScarlet (red) and Hsf1-YFP (green) along the dashed lines in the merged images in (F).

### Sis1 localizes to the nucleolar periphery and cytosolic foci during heat shock

To investigate the localization of Sis1 during heat shock, we tagged Sis1 with YFP to enable live cell fluorescence imaging over a heat shock time course. In the Sis1-YFP cells, we tagged the disaggregase chaperone Hsp104 with mKate. Hsp104 is among the strongest Sis1 interactors (Figure 3A) and is known to form cytosolic foci during heat shock. Hsp104 foci are an indicator that a given cell is stressed. Prior to heat shock, Sis1-YFP was concentrated in the nucleus in all cells but also showed diffuse cytosolic signal along with a few cytosolic foci that colocalized with Hsp104-mKate (Figure 4A). Rapidly upon heat shock, Sis1-YFP re-localized to form a ring in the nucleus and, in concert with Hsp104-mKate, coalesced into multiple cytosolic puncta (Figure 4A, Movie S1).

**Figure 4.**
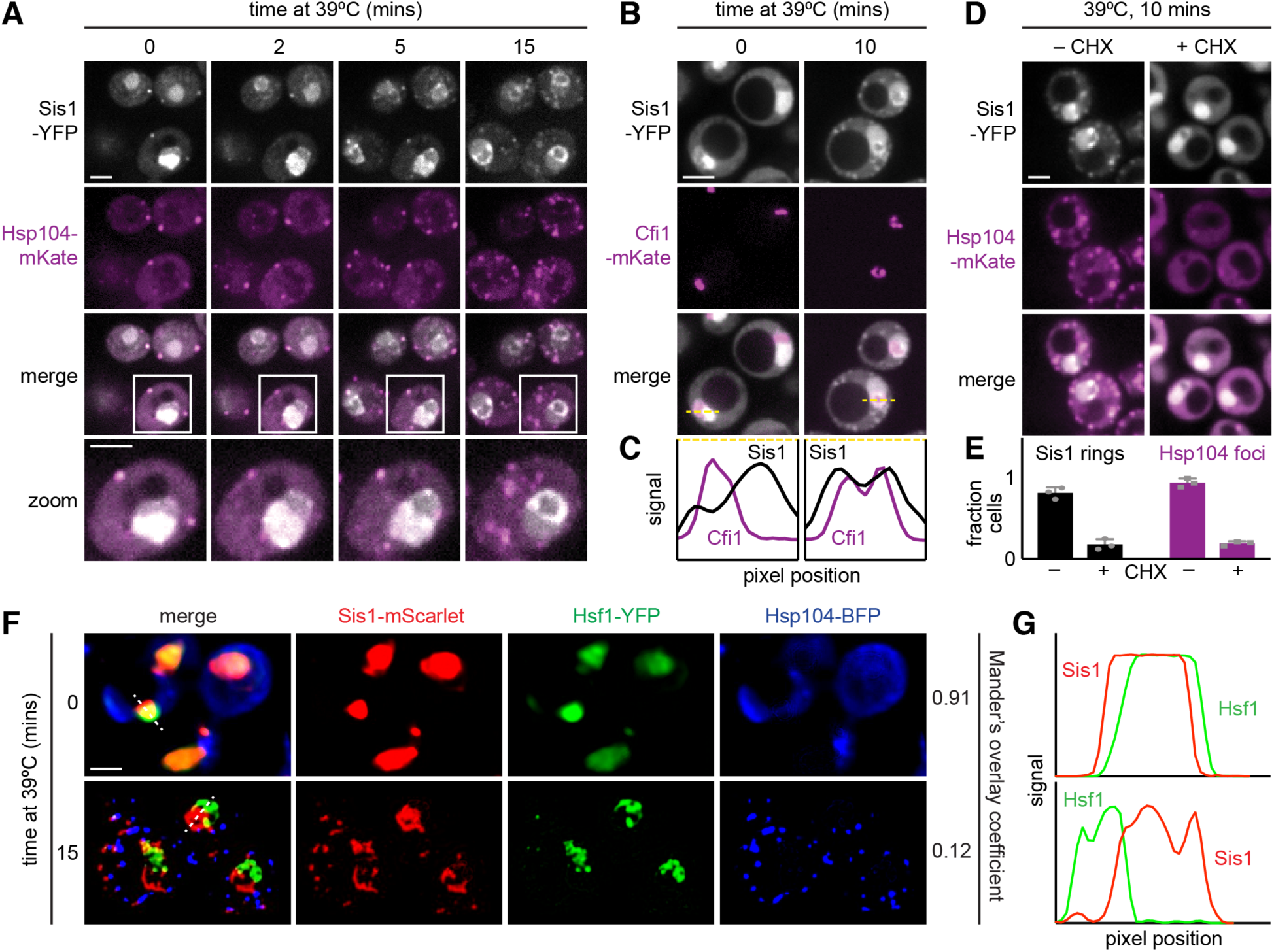
The Sis1 interactome during heat shock. **A)** Anti-FLAG IPs were performed from an untagged strain and a strain expressing Sis1-3xFLAG both heat-shocked for 15 minutes at 30°C. The significance over background is plotted as a function of the apparent stoichiometry (stoichiometry_app_) – the ratio of background-subtracted quantitative value of each of the 192 interacting proteins to the value of Sis1. Proteins are color-coded to match the categories in (C). **B)** Gene ontology associations enriched among the Sis1 interacters during heat shock. **C)** Sis1 interactors grouped by functional category. Ribosomal proteins are abbreviated with their subunit designation (L or S) and their identifier. Proteins in gray are all those that do not belong to the other groups, enriched for highly expressed cytosolic enzymes.

The nuclear ring encompassed a sub-region of the nucleus that was relatively depleted of Sis1-YFP under nonstress conditions. Since Sis1 interacted with a set of nucleolar proteins (Figure 3C), we hypothesized this nuclear sub-region to be the nucleolus, especially given its crescent-shaped morphology in unstressed cells (Figure 4A) (*38*). To monitor Sis1 localization relative to the nucleolus, we imaged Sis1-YFP in a strain in which we tagged the nucleolar resident protein Cfi1 with mKate. Under nonstress conditions, Cfi1-mKate localized to the sub-region of the nucleus we expected. Following heat shock, Sis1-YFP formed a ring around Cfi1-mKate (Figure 4B). Fluorescence intensity line scans showed that the signal peaks for Cfi1-mKate and Sis1-YFP are adjacent prior to heat shock; upon heat shock, the Cfi1-mKate signal becomes surrounded by the Sis1-YFP signal (Figure 4C). Thus, Sis1 appears to encircle the nucleolus during heat shock.

The nucleolus is the site of ribosome biogenesis, and nascent ribosomal proteins traffic from the cytosol where they are synthesized to the nucleolus where they are incorporated with rRNA into large and small ribosomal subunits. Unincorporated “orphan” ribosomal proteins (oRPs) are known to be potent activators of Hsf1 in the absence of stress (*39, 40*), and ribosomal proteins constituted greater than 30% (59/192) of the Sis1-interacting proteins (Table S3). Ongoing protein synthesis has been shown to be required for Hsp104 to form foci and for full Hsf1 activation during heat shock (*15, 39*). To block translation in general and the production of ribosomal proteins in particular, we added cycloheximide prior to heat shocking cells expressing Sis1-YFP and Hsp104-mKate. In the presence of cycloheximide, we observed strong reduction in both Hsp104-mKate cytosolic foci and in Sis1 nucleolar ring formation (Figure 4D, E). This suggests that ongoing translation is required to trigger Sis1 re-localization during heat shock, and is consistent with a role for nascent ribosomal proteins in Sis1 localization to the nucleolus. However, whether oRPs are the biochemical entities that recruit Sis1 to the nucleolus mains to be determined.

Regardless of the species involved, an implication of Sis1 re-localization to the nucleolar periphery is that it may be depleted from the nucleoplasm where Hsf1 resides. To determine if Sis1 localizes away from Hsf1 during heat shock, we tagged Sis1 with mScarlet in a strain expressing Hsf1-YFP and Hsp104-BFP. Under nonstress conditions, Sis1-mScarlet and Hsf1-YFP both show diffuse nuclear localization patterns, and Hsp104-BFP is diffuse in the cytosol (Figure 4F). Quantification revealed >90% signal overlap between Sis1-mScarlet and Hsf1-YFP (Mander’s overlap coefficient (MOC) = 0.91) (*41*). Upon heat shock, Sis1-mScarlet and Hsp104-BFP formed puncta in the cytosol, Sis1-mScarlet formed a subnuclear ring and Hsf1-YFP formed subnuclear clusters (Figure 4F). Despite both being in the nucleus, Sis1-mScarlet and Hsf1-YFP showed nearly mutually exclusive spatial patterns and >7-fold reduction in signal overlap during heat shock (MOC = 0.12) (Figure 4G). Taken together, these data demonstrate that Sis1 localizes to a perinucleolar ring and cytosolic foci during heat shock – and away from Hsf1 – provided that cells are actively translating.

### Sis1 colocalizes with the proteasome and RQC in an interconnected network

In addition to nucleolar factors, the IP/MS dataset revealed Sis1 interactions with cytosolic and ER factors (Figure 3C). To resolve the subcellular organization of Sis1 with high spatial resolution, we employed lattice light sheet imaging and deconvolution to generate 3D reconstructions of cells under nonstress and heat shock conditions (*42*). We chose to focus on the proteasome and the ribosome-associated quality control complex to investigate how these proteostasis modules interact with Sis1. We constructed tri-color strains expressing Sis1-YFP, Hsp104-BFP and proteasome or RQC factors tagged with mScarlet. Prior to heat shock, the proteasome subunit Rpn1-mScarlet localized diffusely to the nucleus and showed a high level of overlap with Sis1-YFP (MOC = 0.63) (Figure 5A). Upon heat shock, Sis1-YFP and Rpn1-mScarlet colocalized in a ring and a network of foci (MOC = 0.82) (Figure 5A, Movie S2). Thus, Rpn1 appears to move with Sis1 to a nucleolar ring and cytosolic foci. The presence of the Rpn1-mScarlet at these sites suggests that Sis1 is localizing to subcellular sites of proteasomal degradation.

**Figure 5.**
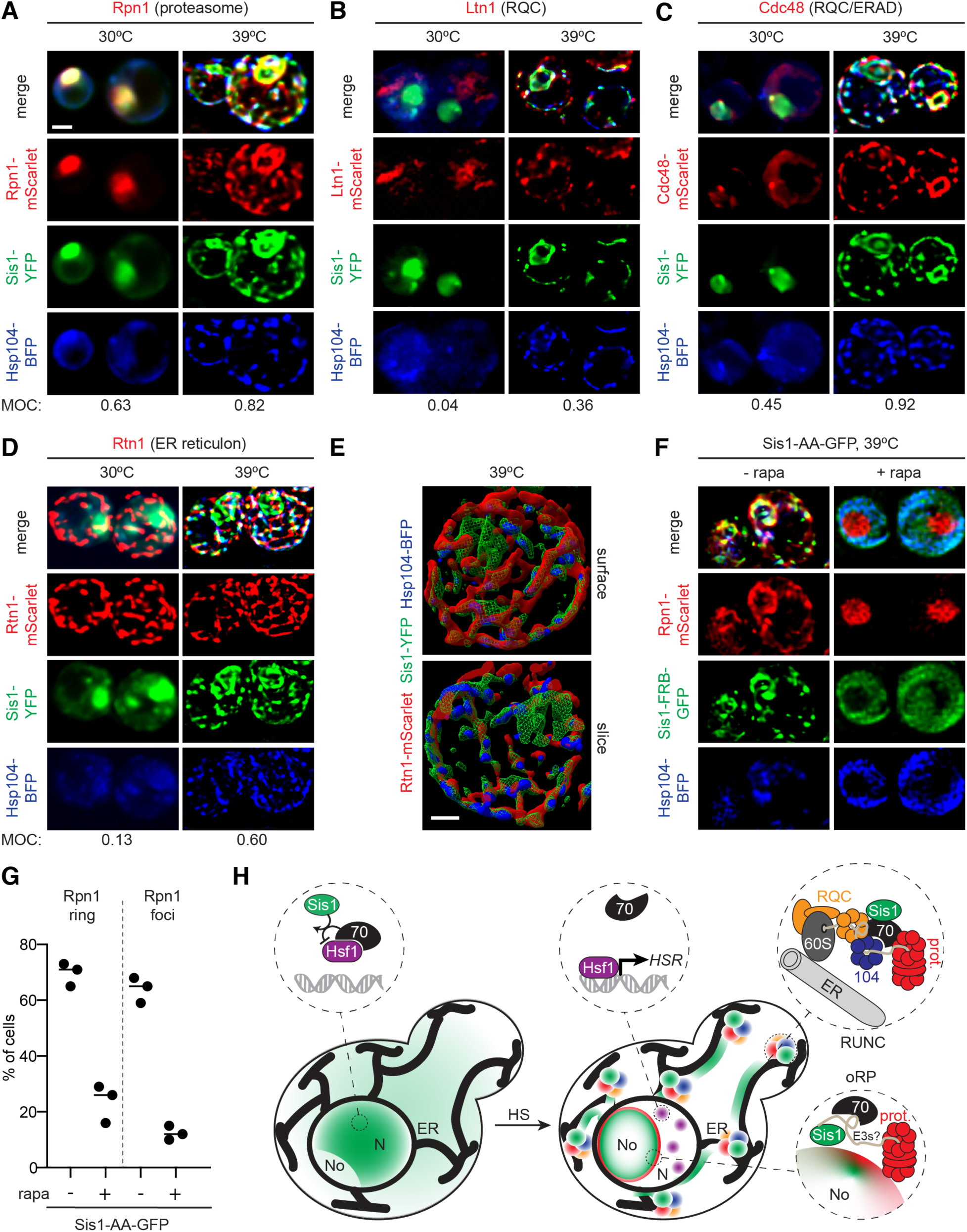
Sis1 forms an interconnected spatial network with other proteostasis factors during heat shock. **A-D)** Scale bar is 2 µm. Deconvolved lattice light sheet 3D reconstructions of live cells under nonstress conditions and 15 minutes of heat shock at 39°C expressing endogenously-tagged Sis1-YFP and Hsp104-BFP with: **A)** Rpn1-mScarlet, a subunit of the proteasome. **B)** Ltn1-mScarlet, a subunit of the ribosome quality control complex (RQC). **C)** Cdc48-mScarlet, a component of ERAD and RQC. **D)** Rtn1-mScarlet, a reticulon protein component of the ER membrane. **E)** Top: Space-filling 3D surface rendering of a cell at 39°C showing the association of Sis1-YFP (green) and Hsp104-BFP (blue) with the inside surface of the reticulated ER as marked by Rtn1-mScarlet (red). Bottom: coronal slice at the midpoint of the cell above. The Sis1-YFP subnuclear ring (green) can be shown and is disconnected from the cytosolic Rtn1 network. Scale bar is 1 µm. **F)** Anchor away of Sis1 precludes formation the spatial proteostasis network. Cells expressing Hsp104-BFP and Rpn1-mScarlet to mark the proteasome in a Sis1-AA background with Sis1-AA-GFP were imaged by a lattice light sheet microscope following 15 minutes of heat shock at 39°C following no pretreatment or pretreatment with rapamycin to anchor away Sis1. **G)** The fraction of cells showing Rpn1 subnuclear rings and cytosolic foci was quantified during heat shock in cells with no pretreatment or rapamycin pretreatment to anchor away Sis1. Experiments were performed in triplicate with >20 cells per replicate. **H)** Cartoon model of how the heat shock-dependent spatial reorganization of the protein homeostasis network is coupled to the regulation of the heat shock response. Left: in the absence of stress, Sis1 is diffuse throughout the cell and concentrated in the nucleus. In the nucleus, it activates Hsp70 to repress Hsf1. Right: Upon heat shock, Sis1 re-localizes to the periphery of the nucleolus and the surface of the ER in the cytosol. In the nucleoplasm, Hsf1 is now free of Hsp70 and can cluster and activate the heat shock transcriptional response. Sis1 recruits the proteasome to colocalize at the nucleolar periphery. On the ER, Sis1 interacts with stalled ribosomes with nascent chains, the ribosome quality control complex (RQC), Hsp104 and the proteasome.

We next investigated the localization of RQC components Ltn1 and Cdc48. RQC is a protein complex that recognizes stalled ribosomes and promotes proteasomal degradation of nascent chains (*20*). Ltn1 is an E3 ligase that modifies nascent chains with ubiquitin, and Cdc48 is a AAA ATPase that mediates the handoff of ubiquitylated nascent chains to the proteasome. Ltn1-mScarlet showed patchy cytosolic staining in unstressed cells and very little colocalization with Sis1-YFP (MOC = 0.04) (Figure 5B). However, upon heat shock, Ltn1-mScarlet increased its overlap with Sis1-YFP 9-fold (MOC = 0.36) and formed foci at the nuclear periphery and cell cortex, some of which colocalized with Sis1-YFP (Figure 5B). In the absence of stress, Cdc48-mScarlet displayed diffuse cytosolic signal and formed foci at the nuclear periphery that overlapped with Sis1-YFP (MOC = 0.45) (Figure 5C). Heat shock triggered Cdc48-mScarlet re-localization to puncta at the cell cortex and nuclear periphery that largely overlapped with Sis1-YFP (MOC = 0.92) (Figure 5C). The difference in localization patterns between Ltn1 and Cdc48 can be attributed to the fact that, in addition to RQC, Cdc48 also participates in ER-associated degradation (ERAD) (*43*). Together these images show that Sis1 colocalizes with RQC during heat shock.

Beyond colocalization during heat shock, the 3D reconstructions of the lattice light sheet images revealed that Sis1-YFP, Hsp104-BFP, Rpn1-mScarlet, Ltn1-mScarlet and Cdc48-mScarlet form a semi-contiguous network (Figure 5A-C). Sis1-YFP and Rpn1-mScarlet form an overlapping and highly ordered network that connects the nucleolar ring with a series of cytosolic foci (Figure S4A, B, Movie S2). Hsp104 is excluded from the nucleolar ring but colocalizes with Sis1 and Rpn1 in the cytosolic network. Hsp90 (Hsc82-mScarlet) does not appear to participate in the network nor in any other Sis1-containing structures (Figure S4C).

The ER has been previously implicated as an organizational factor for the spatial arrangement of protein quality control factors, including Hsp104 (*27, 44*). Moreover, we found that the ER structural protein Rtn1 co-precipitated with Sis1-3xFLAG during heat shock. To test if the Sis1 cytosolic network forms in proximity to the ER, we imaged Rtn1-mScarlet in cells expressing Sis1-YFP and Hsp104-BFP. Indeed, Sis1-YFP increased its signal overlap with Rtn1-mScarlet from MOC = 0.13 under nonstress conditions to MOC = 0.60 during heat shock (Figure 5D), suggesting increased association with the ER. Space-filling 3D cell projections reveal the orientation of the interaction network, with Rtn1 toward the periphery and Sis1 and Hsp104 lining the interior (Figure 5E). The imaging data suggest that Sis1 forms a highly connected network with the proteasome and RQC on the surface of the ER.

Lastly, we tested if Sis1 is required for the proteasome to re-localize to the nucleolar ring during heat shock. To this end we anchored away Sis1 prior to heat shocking cells and monitored localization of Rpn1-mScarlet and Hsp104-BFP. Following Sis1 anchor away, we found that Rpn1-mScarlet remained nuclear during heat shock and failed to form a nucleolar ring (Figure 6F, G). The short period of treatment (15 minutes) makes it unlikely that Sis1 anchor away pre-adapted cells to heat shock and thus precluded the stress. Rather, Sis1 appears to be necessary to recruit the proteasome. This implies that Sis1 mediates proteasomal degradation of chaperone clients on the surface of the nucleolus.

## DISCUSSION

Here we showed that localization of the J-protein Sis1 is a key determinant of Hsf1 activity and transcriptional induction of the heat shock response. Artificial nuclear depletion of Sis1 triggers Hsf1 dissociation from Hsp70 and specifically activates the HSR. During physiological heat shock, Sis1 leaves the nucleoplasm and re-localizes to a subnuclear ring and cytosolic foci. The Sis1 subnuclear ring surrounds the nucleolus and recruits the proteasome. The cytosolic foci form a semi-contiguous network on the surface of the ER that contains the disaggregase Hsp104, the proteasome and RQC. By spatially collaborating with proteostasis machinery at the surfaces of the nucleolus and ER during heat shock, Sis1 re-localizes away from Hsf1 and thereby ceases to promote Hsp70-mediated Hsf1 repression. Thus, Sis1 couples the spatial remodeling of the proteostasis machinery to regulation of the heat shock response.

The role of Sis1 in promoting Hsp70-mediated repression of Hsf1 addresses a conceptual challenge inherent in the “chaperone titration model” of Hsf1 regulation. How can cytosolic chaperone clients produced by heat shock titrate away Hsp70 in the nucleus to enable Hsf1 activation in less than one minute? This challenge is compounded by the fact that Hsp70 outnumbers Hsf1 by three orders of magnitude in the cell (*25*). Sis1 helps to resolve this conundrum. First, Sis1 is expressed in only 10-fold excess over Hsf1 (*25*). Second, the requirement for Sis1 – a J-protein that activates Hsp70 to hydrolyze ATP – implies that the relevant regulatory species is not the total cellular pool of Hsp70 but rather the smaller pool of nuclear localized ADP-bound Hsp70. As Sis1 moves away from the nucleoplasm during heat shock, the local concentration of Hsp70-ADP near Hsf1 would be expected to drop, thereby de-repressing Hsf1. Complementary to our results, Sis1 was recently identified as the strongest Hsf1 repressor in the genome in a CRISPRi-based screen for Hsf1 regulators (*45*).

Sis1 re-localization to the surface of the nucleolus during heat shock is consistent with an emerging view of the nucleolus as a protein quality control compartment during stress (*46*). While such partitioning may be a driving force for Sis1 to move to the nucleolus, our observation that blocking translation with cycloheximide reduces Sis1 spatial re-localization during heat shock suggests nascent proteins are also drivers (Figure 3D, E). The nucleolus is the site of ribosome biogenesis, and yeast cells must generate more than 10^5^ nascent ribosomal proteins per minute to support cell division (*47*). Until they are embedded in the ribosome, ribosomal proteins are unstable and aggregation-prone (*39, 40*). Moreover, ribosomal proteins contain many positively charged and hydrophobic residues – precisely the amino acids enriched in Hsp70 substrate binding sites (*48*). Indeed, “orphan ribosomal proteins” (oRPs) have been shown to be potent Hsf1 activators in the absence of other stresses (*39, 40*). Our observation that Sis1 and the proteasome localize to the surface of the nucleolus during heat shock suggests that oRPs may recruit Sis1 to activate Hsp70-mediated proteasomal degradation (Figure 5E).

In the cytosol, Sis1 also displays stress-dependent re-localization. 3D reconstructions revealed that – rather than forming a series of discrete foci – Sis1 and Hsp104 form a semi-contiguous network along with the proteasome and components of the RQC (Figure 5A-C, S4A, B). In addition to these proteostasis factors, we identified an interaction between Sis1 and the ER reticulon protein Rtn1 during heat shock (Figure 4A), in agreement with reports that aggregates of Hsp104 and model misfolded proteins are scaffolded by the ER (*27, 44*). Indeed, Sis1, Hsp104, the proteasome and RQC form a network that largely overlaps with the reticulated ER as marked by Rtn1 (Figure 5D, E). We propose that this reticulum of proteostasis factors serves to organize chaperones in two dimensions to triage “ribosomes with unstable nascent chains” (RUNCs) and allow rapid diffusion of degradation machinery (Figure 5H). This reorganization of Sis1 and other proteostasis components during stress demonstrates that, in addition to being a network of co-regulated genes, the proteostasis network forms a dynamic and spatially organized physical structure. Two-dimensional localization of Sis1 on the surface of the nucleolus and ER may enable rapid coordination of the proteostasis machinery in the nucleus and cytosol with the HSR.

## ACKNOWLEDGEMENTS

We thank H. Yoo and D. A. Drummond for providing important insights into the biochemical role of Sis1. We thank J. Greenberg, E. Ferguson, G. Bushkin, and the other members of the Pincus lab for insightful discussions and comments on the manuscript. We are grateful to T. Volkert and S. Gupta at the Whitehead Genome Technology Core and E. Spooner at the Whitehead Proteomics Core Facility for technical assistance. The initial phase of this work was funded by an Early Independence Award from the NIH Office of the Director (DP5 OD017941 to D.P.), while the later phase was funded by institutional startup provided by the Department of Molecular Genetics and Cell Biology in the Biological Sciences Division at the University of Chicago to D.P.

## AUTHOR CONRIBUTIONS

Conceptualization: D.P.; Investigation: Z.A.F, A.A., J.K., X.Z., D.W., V.P.B., D.P.; Methodology: V.P.B., D.W., D.P.; Formal Analysis: A.S.; Writing – Original Draft: D.P.; Writing – Review & Editing: all authors; Visualization: D.P.; Supervision: S.J.K, D.P.

## METHODS

### Strain construction and cell growth

Yeast strains used in this study are listed in Table S1. All strains are derivatives of W303 and fluorescent protein and epitope tags are integrated into the genome. Cells were cultured in SDC media for confocal imaging and flow cytometry experiments, and in SDC-riboflavin and folic acid for low autofluorescence for lattice light sheet imaging. Cells were cultured in YPD media for immunoprecipitation and RNA seq experiments. Heat shocks were performed at 39°C. Anchor away experiments were conducted using 1 µM rapamycin.

### HSE-YFP flow cytometry assays

Reporter levels in untreated, endpoint and time course assays from heat-shocked and rapamycin-treated cells were measured using a BD Fortessa flow cytometer in the Whitehead Flow Cytometry Facility or the University of Chicago Cytometry and Antibody Technology Facility equipped with a high throughput sampler and analyzed using FlowJo as previously described (*16*).

### RNA-seq sample prep and analysis

Total RNA was purified from yeast as described (*36*) and polyA+ RNA seq libraries were constructed using the NEB Next Ultra RNA kit. Sequencing was performed on an Illumina Hi-Seq 2500 at the Whitehead Institute Genome Technology Core Facility and reads were aligned and quantified as previously described (*36*). Raw sequence files and processed data have been deposited at the Gene Expression Omnibus (GEO, accession number GSE145936).

### Immunoprecipitation

Hsf1-3xFLAG-V5 was serially immunoprecipitated as previously described (*16*) for IP/Western analysis. For IP/MS of Sis1-3xFLAG-V5 and Hsf1-3xFLAG-V5, the protocol was modified to only perform the anti-FLAG IP using a short incubation of the anti-FLAG beads with total lysate for 15 minutes prior to washing and eluting with 3xFLAG peptide. The short incubation was designed to increase the likelihood of capturing transient interactions while reducing nonspecific interactions. All IP experiments were performed in biological triplicate. Triplicate IPs were also performed on an untagged strain exposed to the same conditions to subtract background and calculate significance.

### Mass spectrometry

MS analysis was performed at the Whitehead Proteomics core for the 15 minute heat shock Sis1-3xFLAG-V5 as described previously (*16*). For the Sis1-3xFLAG-V5 untreated samples and the Hsf1-3xFLAG-V5 samples, MS analysis was performed as follows:

#### In-Solution Trypsin Digestion

30 µl of eluate was in-solution digested with Trypsin by first reducing in 50mM ABC with 6 µl Rapigest surfactant (Waters) and 10% 200 mM TCEP. Alkylated with 50mM Iodoacetamide (33 µl) in dark 30 minutes at RT. Digested 1:50 v/v Trypsin (Promega) at 37°C o/n. Detergent was removed with 1 µl TFA at 37C for 45 minutes. Digested peptides were cleaned up on a C18 column (Pierce), speed vac’d and sent for LC-MS/MS to the Proteomics Core at Mayo Clinic.

#### HPLC for mass spectrometry

All samples were resuspended in Burdick & Jackson HPLC-grade water containing 0.2% formic acid (Fluka), 0.1% TFA (Pierce), and 0.002% Zwittergent 3-16 (Calbiochem), a sulfobetaine detergent that contributes the following distinct peaks at the end of chromatograms: MH+ at 392, and in-source dimer [2M + H+] at 783, and some minor impurities of Zwittergent 3-12 seen as MH+ at 336. The peptide samples were loaded to a 0.25 μl C8 OptiPak trapping cartridge custom-packed with Michrom Magic (Optimize Technologies) C8, washed, then switched in-line with a 20 cm by 75 μm C18 packed spray tip nano column packed with Michrom Magic C18AQ, for a 2-step gradient. Mobile phase A was water/acetonitrile/formic acid (98/2/0.2) and mobile phase B was acetonitrile/isopropanol/water/formic acid (80/10/10/0.2). Using a flow rate of 350 nl/min, a 90 minute, 2-step LC gradient was run from 5% B to 50% B in 60 minutes, followed by 50%-95% B over the next 10 min, hold 10 min at 95% B, back to starting conditions and re-equilibrated.

#### LC-MS/MS data acquisition and analysis

The samples were analyzed via data-dependent electrospray tandem mass spectrometry (LC-MS/MS) on a Thermo Q-Exactive Orbitrap mass spectrometer, using a 70,000 RP survey scan in profile mode, m/z 360-2000 Da, with lockmasses, followed by 20 HCD fragmentation scans at 17,500 resolution on doubly and triply charged precursors. Single charged ions were excluded, and ions selected for MS/MS were placed on an exclusion list for 60 s. An inclusion list was utilized consisting of expected prototypic peptide ions in the 2+ and 3+ charge state for the yeast proteins ySIS1 and yHSF1 using in-house software. All LC-MS/MS *.raw Data files were analyzed with MaxQuant version 1.5.2.8, searching against the SPROT Yeast database (Downloaded 5/23/2019 with isoforms, 12154 entries) *.fasta sequence, using the following criteria: LFQ was selected for Quantitation with a minimum of 1 high confidence peptide to assign LFQ Intensities. Trypsin was selected as the protease with maximum missing cleavage set to 2. Carbamiodomethyl (C) was selected as a fixed modification. Variable modifications were set to Oxidization (M), Formylation (N-term), Deamidation (NQ), and Phospo (STY).

Orbitrap mass spectrometer was selected using an MS error of 20 ppm and a MS/MS error of 0.5 Da. 1% FDR cutoff was selected for peptide, protein, and site identifications. Ratios were reported based on the LFQ Intensities of protein peak areas determined by MaxQuant (version 1.5.2.8) and reported in the proteinGroups.txt. The proteingroups.txt file was processed in Perseus (version 1.6.7). Proteins were removed from this results file if they were flagged by MaxQuant as “Contaminants”, “Reverse” or “Only identified by site”. Three biological replicates were performed. Samples were filtered to require hits to have been seen in at least two replicates per condition. LFQ peak intensities Log2 transformed and median normalized and missing values were imputed via default settings in Perseus.

### Mathematical modeling

To model the Sis1-Hsp70-Hsf1 circuit we have expanded the differential equation model from Zheng et al. eLife 2016 to include the effects of Sis1. The model consists of five different protein species (Hsf1, Hsp70, client protein (UP), Sis1, reporter protein YFP) and two protein complexes:

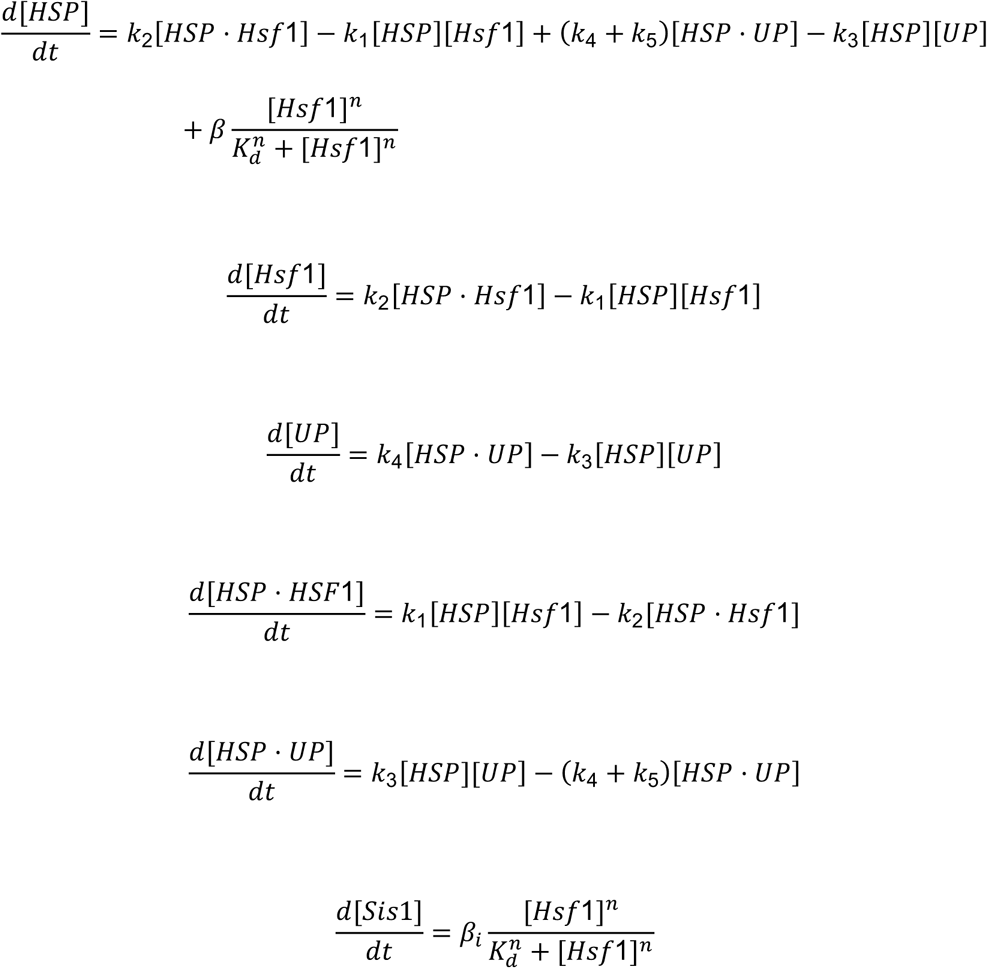

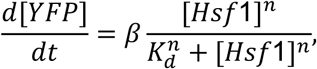

where [] denotes the nuclear concentration of respective species. We refer the reader to Zheng et al. eLife 2016 for modeling details, assumption, and parameter values. The rate *k*_1_ = 184 *min*^−1^ *a. u*. ^−1^ denotes the binding of Hsp70 to Hsf1 to create an inactive complex HSP· Hsf1, and the complex dissociates with rate *k*_2_. Sis1 enhances the repression of Hsf1 by Hsp70, and we phenomenologically capture this by assuming that *k*_2_ is dependent on Sis1 levels as per the following Hill equation

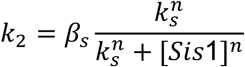

with *n* = 3, *β*_*s*_ = 1.12 *min*^−1^ and *k*_*s*_ = 1.54 *a. u*. The rate *k*_3_ = 136 *min*^−1^ *a. u*. ^−1^ is the binding of Hsp70 to client proteins to create the complex HSP · UP that dissociates with rate *k*_4_ = 0.06 *min*^−1^. The degradation of UP by Hsp70 is captured via the rate *k*_5_ = 10^−5^ *min*^−1^. The activation of both YFP and Hsp70 by Hsf1 is modeled by a Hill equation with *n* = 3, *β* = 1.8 *min*^−1^ and *k*_*d*_ = 0.0057 *a. u*. The activation of Sis1 is similarly modeled but with activation rate *β*_9_ that depends on the nuclear export signal.

The above differential equation model was run with the following initial values (in a.u.) at time *t* = 0

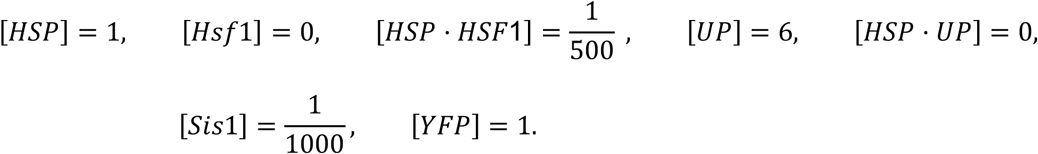

The effect of nuclear localization signals was modeled by changing a single parameter *β*_*i*_ with *β*_*i*_ = 1.86 *min*^−1^ for WT and *β*_*i*_ = 4.1 *min*^−1^ for NLS.

### Confocal imaging

Confocal imaging was performed at the Nikon Imaging Center at the Whitehead Institute for Biomedical research using the Andor spinning disc system as previously described (*23*). Heat shock was performed on live cells using an objective heater as described (*24*).

### Lattice light sheet imaging and analysis

Lattice light sheet imaging was performed at the University of Chicago Integrated Light Microscopy Core using a phase-2 system designed by Intelligent Imaging Innovations (3i) and run in SlideBook 6.0 software (3i). The design is a modification of the original (*42*), with greater automation and stability. Optics were aligned daily, and bead point-spread function (PSFs) collected prior to imaging cells. The imaging camera was a Hamamatsu Fusion chilled sCMOS run at default speed/quality. The annulus mask was set for a 20 μm beam length (outer NA 0.55 inner NA 0.493), 400 nm thickness, with dither at 9 μm. Laser intensities were set to balance signal and bleaching rates. Temperature was controlled by a built-in Peltier device. Sample scan image stacks were deskewed in SlideBook, and the tif series were processed using NIH ImageJ [Fiji version]. GPU-based Richardson-Lucy deconvolution used measured PSFs or theoretical PSFs via Brian Northan’s “OPS” implementation (https://github.com/imagej/ops-experiments). Reconstructions and movies were assembled using ClearVolume (*49*).

## FIGURE LEGENDS

**Figure S1.**
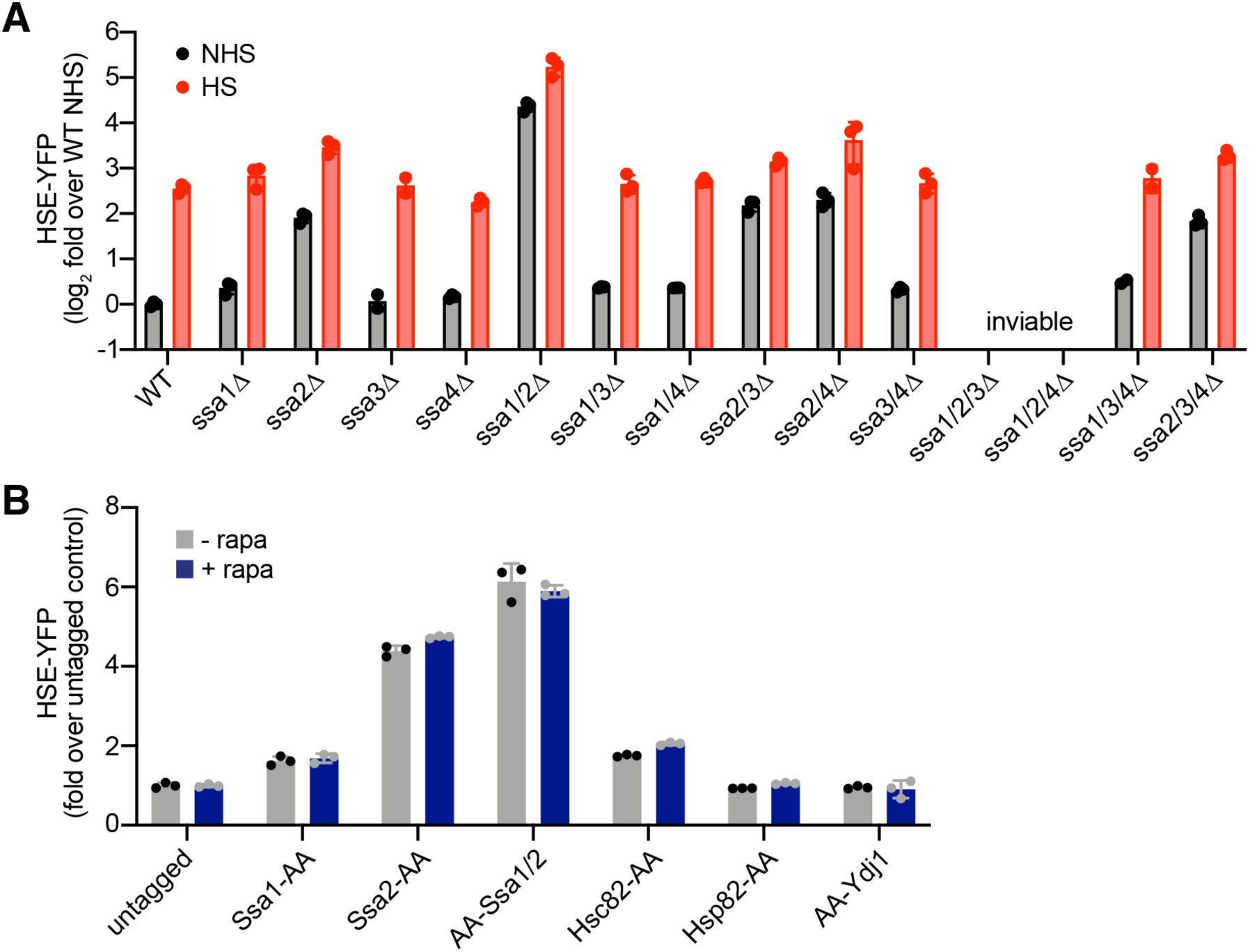
Hsf1 activity in Hsp70 mutants and chaperone anchor away strains. **A)** Heat shock element (HSE)-YFP reporter assay for Hsf1 activity in Hsp70 deletion strains. Cells were left untreated or heat shocked at 39°C for four hours and YFP levels were measured by flow cytometry and normalized to untreated wild type. Three biological replicates are shown, along with the mean and standard deviation. **B)** HSE-YFP reporter assay of anchor away strains in the presence and absence of rapamycin normalized to the untagged anchor away parent strain. Cells were treated with 1 µM rapamycin for 8 hours before measuring the reporter. These data suggest that C-terminal tagging of Ssa1 and Ssa2 compromises their function nearly as severely as knocking them out. Consistent with this interpretation, individual AA-tagging of Ssa1 and Ssa2 resulted in mild and moderate increases HSE-YFP levels, respectively, akin to their respective deletions. N-terminal AA tagging of Ssa2 likewise resulted in a constitutive increase in HSE-YFP signal.

**Figure S2.**
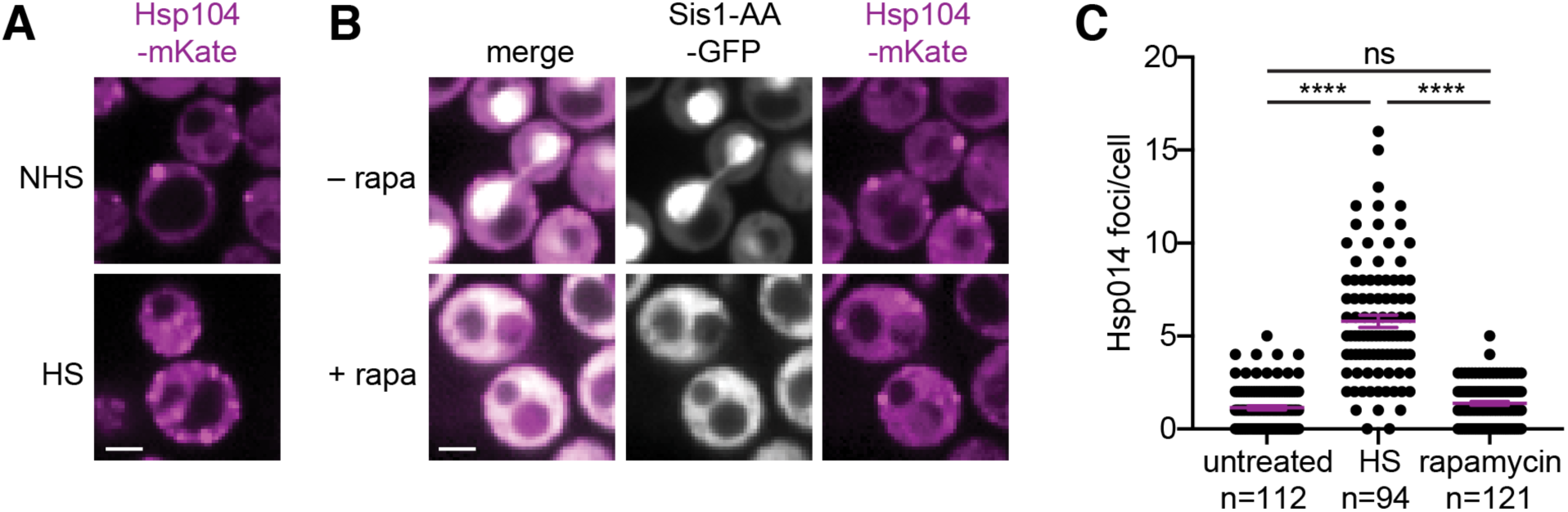
Anchor away of Sis1 does not trigger proteostasis collapse. **A)** Cells expressing Hsp104-mKate were imaged by spinning disc confocal microscopy under non heat shock (NHS) and following 15 minutes of heat shock (HS) at 39°C. Scale bar is 2 µm. **B)** Cells expressing Hsp104-mKate and Sis1-AA-GFP in untreated cells and following the addition of rapamycin for 1 hour to anchor away Sis1-AA-GFP. The two dark regions in the rapamycin-treated cells are the nucleus and the vacuole. **C)** Quantification of the number of Hsp104-mKate foci in individual cells either left untreated, following heat shock, or following Sis1 anchor away with rapamycin.

**Figure S3.**
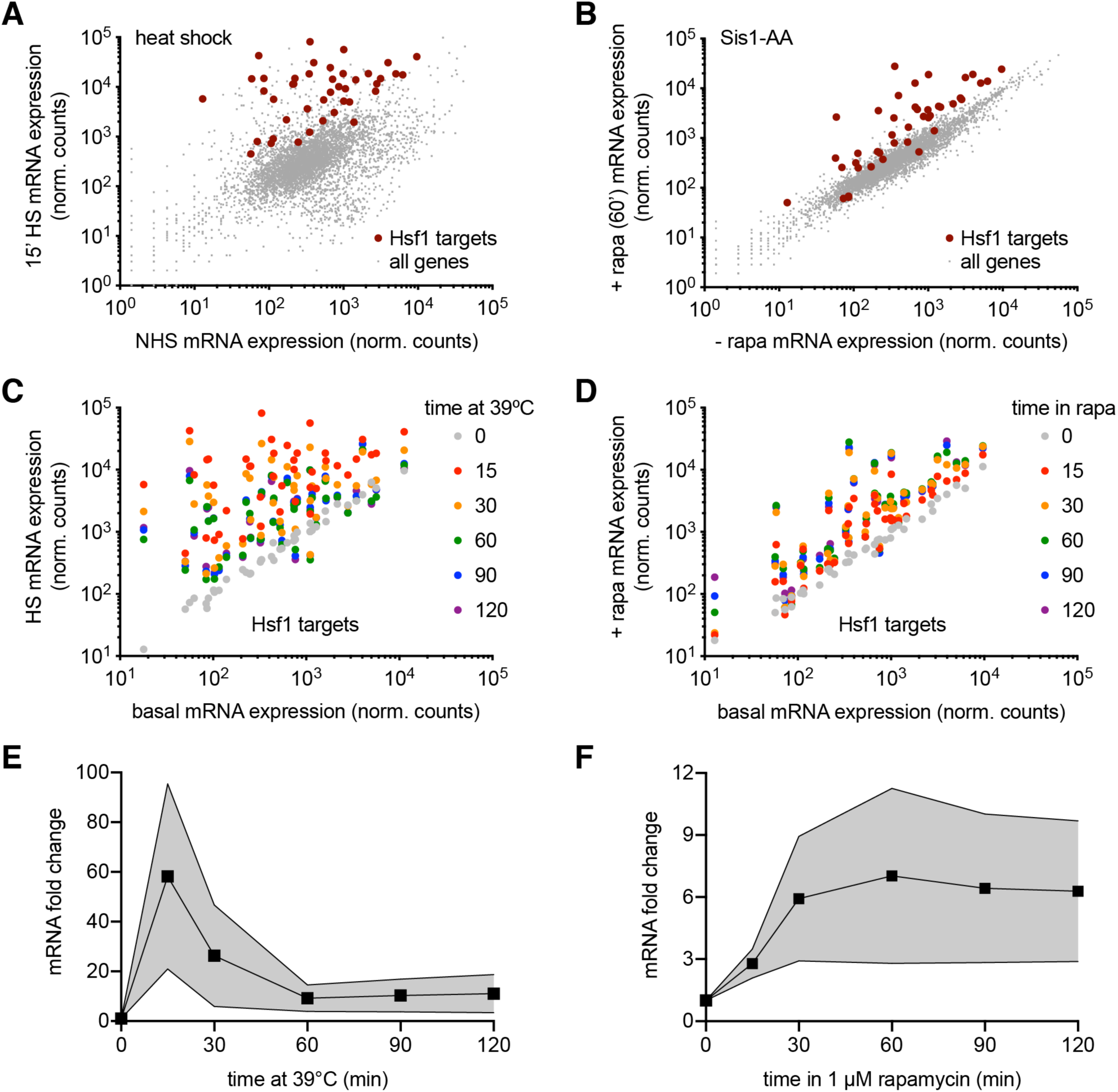
Comparative transcriptomics following heat shock and Sis1 anchor away. **A)** RNA-seq of heat shocked cells (15 minutes at 39°C) versus non-stressed cells. In addition to induction of the Hsf1 target genes (dark red), many other genes are induced and repressed. **B)** RNA-seq of cells with Sis1 anchored away for 60 minutes versus untreated cells. Hsf1 target genes are specifically induced without other changes to the transcriptome. **C)** All 42 Hsf1 target gene expression levels over a heat shock time course. Most genes peak at the 15-minute time point. **D)** All 42 Hsf1 target gene expression levels of a time course following Sis1 anchor away. Levels do not decline over time and peak at later time points for most genes. **E)** The average and standard deviation of the induction of the 42 Hsf1 target genes over a heat shock time course showing the adaptive response. **F)** As in (E) but for the Sis1 anchor away time course. There is no adaptation, suggesting the feedback loop has been severed.

**Figure S4.**
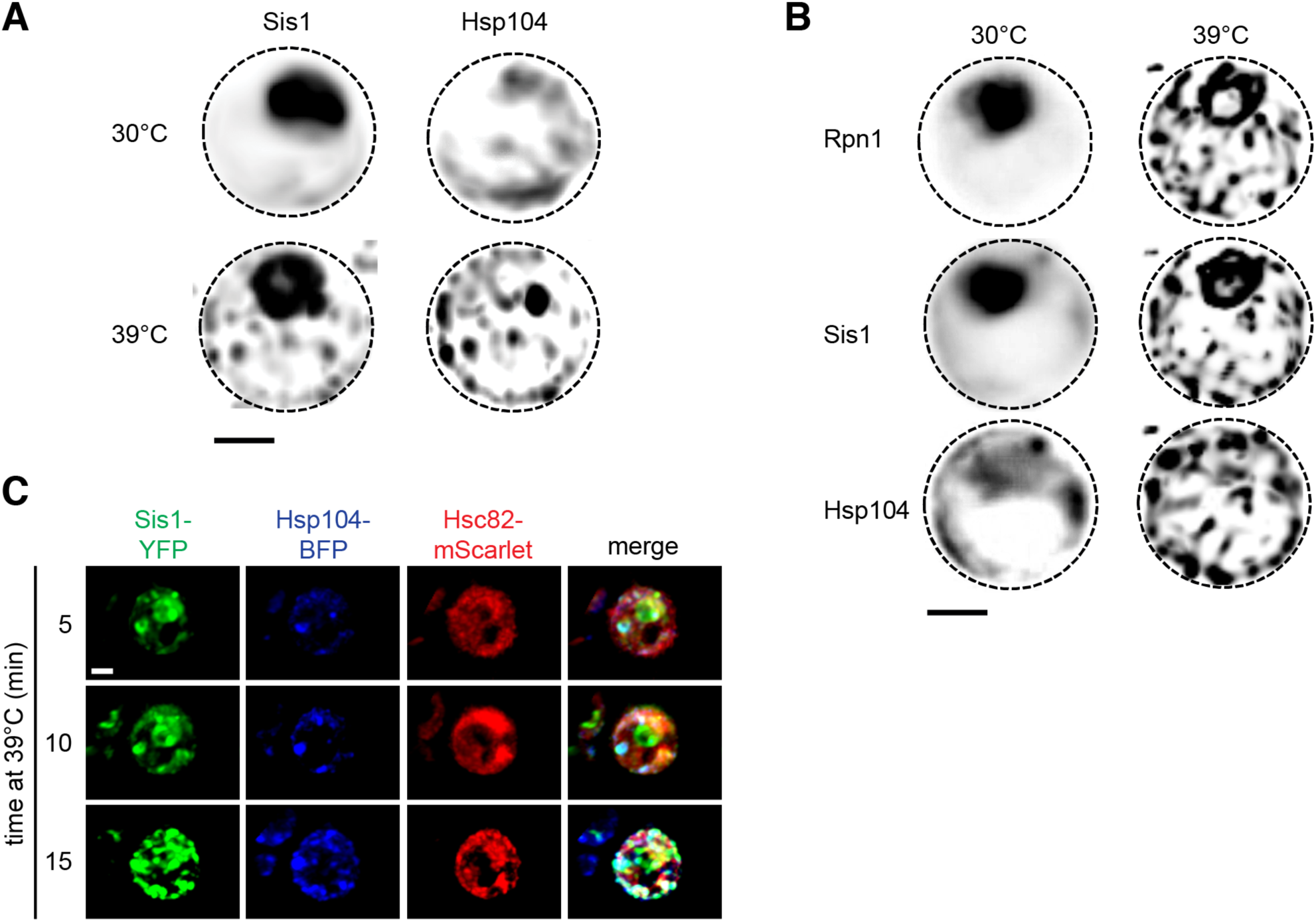
The proteostasis machinery forms a contiguous network on the ER. **A)** Maximum intensity negative projections of Sis1-YFP and Hsp104-BFP signal in cells under nonstress and heat shock conditions (30°C for 15 minutes). The signal is enhanced to show the cytosolic interconnections between the nodes. Scale bar is 2 µm. **B)** As in (A) but for the proteasome component Rpn1. **C)** Heat shock time course of a cell expressing Sis1-YFP, Hsp104-BFP and Hsc82-mScarlet. Hsc82 is the yeast homolog of Hsp90. Hsc82 neither associates with the nucleolar ring nor participates in the cytosolic network. Scale bar is 2 µm.

**Movie S1**.

Live imaging on a spinning disc confocal of cells growing at 25°C expressing Sis1-YFP (white) and Hsp104-mKate (magenta) shifted to 39°C and imaged over time. 45 seconds elapse between frames.

**Movie S2**.

3D rotation of a lattice light sheet image of a cell expressing Sis1-YFP (green), Rpn1-mScarlet (red) and Hsp104-BFP (blue) at 39°C for 15 minutes. Rpn1 marks the proteasome.

**Movie S3**.

3D rotation of a lattice light sheet image of a cell expressing Sis1-YFP (green), Cdc48-mScarlet (red) and Hsp104-BFP (blue) at 39°C for 15 minutes. Cdc48 functions in RQC and ERAD.

**Movie S4**.

3D rotation of a lattice light sheet image of a cell expressing Sis1-YFP (green), Rtn1-mScarlet (red) and Hsp104-BFP (blue) heat shocked at 39°C for 15 minutes. This movie shows colocalization of Sis1 and Hsp104 with the reticulated ER during heat shock.

**Table S1**.

Yeast strains used in this study.

**Table S2**.

Hsf1-3xFLAG IP/MS results.

**Table S3**.

Sis1-3xFLAG IP/MS results.

